# TERRA ONTseq: a long read-based sequencing pipeline to study the human telomeric transcriptome

**DOI:** 10.1101/2023.11.30.569384

**Authors:** Joana Rodrigues, Roberta Alfieri, Silvia Bione, Claus M. Azzalin

**Affiliations:** Instituto de Medicina Molecular João Lobo Antunes (iMM), Faculdade de Medicina da Universidade de Lisboa, Lisbon, 1649-028, Portugal; Istituto di Genetica Molecolare Luigi Luca Cavalli-Sforza – Consiglio Nazionale delle Ricerche, Pavia, 27100, Italy; Istituto di Tecnologie Biomediche – Consiglio Nazionale delle Ricerche, Segrate (MI), 20054, Italy

**Keywords:** telomeres, TERRA, transcription, CpG dinucleotide promoters, long read nanopore sequencing

## Abstract

The long noncoding RNA TERRA is transcribed from telomeres in virtually all eukaryotes with linear chromosomes. In humans, TERRA transcription is driven in part by promoters comprising CpG dinucleotide-rich repeats of 29 base pairs (29 bp repeats), believed to be present in half of the subtelomeres. Thus far, TERRA expression has been analyzed mainly using molecular biology-based approaches that only generate partial and somehow biased results. Here, we present a novel experimental pipeline to study human TERRA based on long read sequencing (TERRA ONTseq). By applying TERRA ONTseq to different cell lines, we show that the vast majority of human telomeres produce TERRA and that the cellular levels of TERRA transcripts varies according to their chromosomes of origin. Using TERRA ONTseq, we also identified regions containing TERRA transcription start sites (TSSs) in more than half of human subtelomeres. TERRA TSS regions are generally found immediately downstream of 29 bp repeat-related sequences, which appear to be more widespread than previously estimated. Finally, we isolated a novel TERRA promoter from the highly expressed subtelomere of the long arm of chromosome 7. With the development of TERRA ONTseq, we provide a refined picture of human TERRA biogenesis and expression and we equip the scientific community with an invaluable tool for future studies.

## INTRODUCTION

Telomeres are protective protein/nucleic acids structures located at the ends of linear eukaryotic chromosomes. In mammalian cells, telomeric DNA is made up of arrays of the (TTAGGG)n sequence, varying in length roughly from 10 to 15 kilobases (kb) amongst different cell types and different chromosome ends within the same cell (Srinivas et al. 2020). Telomeres are transcribed by DNA-dependent RNA polymerase II (RNAPII) into the evolutionary conserved long non-coding RNA TERRA (Azzalin et al. 2007; Azzalin and Lingner 2008; Schoeftner and Blasco 2008). TERRA transcription starts within subtelomeres and proceeds towards the ends of the chromosomes with the C-rich telomeric strand acting as a RNAPII template. Hence, individual TERRA molecules comprise a sequence specific to the subtelomere of origin followed by stretches of UUAGGG repeats (Azzalin et al. 2007; Schoeftner and Blasco 2008; Porro et al. 2010; Farnung et al. 2012).

In humans, transcription of several TERRA molecules initiates at CpG dinucleotide-rich tandem repeats of 29 base pairs (29 bp repeats), which were reported to be present in about half of the subtelomeres (Nergadze et al. 2009; Feretzaki et al. 2019). CpG dinucleotides within 29 bp repeats are heavily methylated, in particular in telomerase positive cells, which normally present relatively low TERRA levels. Simultaneous deletion of the DNA methyltransferases DNMT1 and DNMT3b in HCT116 cancer cells (DKO cells) fully eliminates CpG methylation within 29 bp repeats, leading to a substantial increase in TERRA levels and more avid RNAPII binding to 29 bp and TTAGGG repeats (Nergadze et al. 2009). Based on these data, we proposed that 29 bp repeat sequences are bona fide TERRA promoters whose activity is epigenetically regulated. Importantly, chromosome ends devoid of 29 bp repeats are also highly transcribed in DKO cells (Feretzaki et al. 2019), suggesting the existence of CpG-containing promoters other than the canonical 29 bp repeats. However, due to the lack of a complete and unambiguous sequence of all human subtelomeres, a gap only recently filled with the Telomere-to-Telomere (T2T) sequence of the human genome (Nurk et al. 2022; Rhie et al. 2023), the exact structure of those promoters is still unknown.

Numerous functions have been assigned to TERRA, including telomerase recruitment to telomeres and activity regulation, heterochromatin establishment and overall telomere integrity maintenance (Rivosecchi and Cusanelli 2023; Zeinoun et al. 2023). TERRA and telomere transcription are also essential to sustain telomere elongation in telomerase negative cancers that maintain telomeres through the Alternative Lengthening of Telomeres (ALT) mechanism (Silva et al. 2021; Bhargava et al. 2022; Silva et al. 2022). In agreement, in ALT cells, 29 bp repeat sequences are hypomethylated and TERRA levels are consistently higher compared to telomerase positive cells (Nergadze et al. 2009; Lovejoy et al. 2012; Arora et al. 2014).

Despite the centrality of TERRA in telomere biology, the analysis of TERRA transcripts remains technically challenging due to their low abundance and highly repetitive nature. Up to this date, TERRA quantifications have been mainly done using reverse transcription-quantitative PCR (RT-qPCR) with subtelomeric oligonucleotides or northern blotting with probes detecting (UUAGGG)n sequences (Azzalin et al. 2007; Porro et al. 2010; Arnoult et al. 2012; Deng et al. 2012; Feretzaki et al. 2019). While both methods provide robust data, they also generate partial results. With RT-qPCR, only a few chosen telomeres are analyzed in each experiment; on the other side, when performing northern blotting to detect UUAGGG repeats, total TERRA from all telomeres is simultaneously visualized but the amount of TERRA produced from each chromosome end cannot be determined. Illumina sequencing of purified TERRA was also performed (Porro et al. 2014). However, human subtelomeres comprise heterogeneously repeated sequences shared by multiple chromosome ends (Kwapisz and Morillon 2020), impeding to unequivocally assign small reads, as the ones generated in Illumina sequencing. Indeed, different results were obtained by independent groups starting from the same sets of TERRA Illumina reads (Porro et al. 2014; Montero et al. 2016). Overall, systematic studies where all TERRA molecules are unambiguously analyzed in a quantitative and qualitative way are lacking.

We have established an Oxford Nanopore Technology (ONT)-based long-read sequencing pipeline to sequence and map the subtelomeric tracts of TERRA molecules purified from human cultured cells (TERRA ONTseq). By applying TERRA ONTseq to different human cell lines and using the latest T2T genome as a reference, we show that TERRA transcripts originate from most chromosome ends although transcript levels varies according to the end of origin. We also show that 29 bp repeat-like sequences are present in the majority of subtelomeres and that TERRA transcription often starts immediately after those repeats. Finally, we isolated a polymorphic CpG-rich TERRA promoter sequence that is located within the highly expressed subtelomere of the long arm of chromosome 7 (7q) 7q subtelomere. TERRA-ONTseq is able to readily generate a qualitative and quantitative picture of the TERRA transcriptome and should become a useful tool to study TERRA biogenesis and functions in physiological and pathological contexts.

## RESULTS AND DISCUSSION

### TERRA ONTseq detects transcripts deriving from a multitude of human chromosome ends

The ONT sequencing technology allows to sequence full-length DNA molecules of up to 100 kb, thus bypassing the need to fragment molecules for library preparation and being ideally suited for the sequencing of repetitive nucleic acids (Kwapisz and Morillon 2020; Warburton and Sebra 2023). We affinity purified TERRA transcripts from nuclear RNA prepared from two telomerase positive (HeLa and HEK293T) and two ALT (U2OS and SAOS2) cell lines using a biotinylated DNA oligonucleotide comprising five CCCTAA repeats, which are complementary to the UUAGGG tracts within TERRA molecules (Supplemental Fig. S1A; Supplemental Table S1). Biotinylated oligonucleotides were 3’ end blocked with dideoxycytosine to avoid interference with the subsequent step of reverse transcription. As shown by northern blotting, purified RNA fractions were largely devoid of the U6 small nuclear RNA, while still containing telomeric UUAGGG repeats (Supplemental Fig. S1B). As quantified by RT-qPCR, relative purification efficiencies of TERRA transcripts varied according to the chromosome end (Supplemental Fig. S1C), possibly due different lengths or sequence conservation of the UUAGGG stretch. Overall, TERRA purification was efficient for all cell lines except for SAOS2 (Supplemental Fig. S1C), which were not used for ONT sequencing.

TERRA-enriched RNA was reverse-transcribed with an oligonucleotide complementary to the UUAGGG-repeats (Supplemental Fig. S1A; Supplemental Table S1). Because the C-rich oligonucleotide can hybridize randomly along the (UUAGGG)n stretch, including in close proximity of the subtelomeric part of the transcript, this step assures that the majority of reads contain subtelomeric sequences and can be mapped. However, our protocol cannot provide information on the length of the UUAGGG tract. The obtained cDNA was Nanopore sequenced and quality score analysis identified reads up to 11375, 12310 and 23869 bp long for U2OS, HEK293T and HeLa, respectively (mean read length: 1064, 866, and 964bp, respectively; median read length: 887, 684 and 779 bp, respectively). Preliminarily alignments of the reads to the build T2T-CHM13v2.0/hs1 of the human genome (Nurk et al. 2022) did not detect any splicing signature (data not shown), which is consistent with previous reports (Porro et al. 2014). Hence, we refined the alignment strategy by generating a subtelomeric reference sequence from the T2T genome comprising 25 kb sequences terminating at the fourth telomeric TTAGGG repeat on each chromosome end. Because all cell lines utilized in this study are from female donors, the recently completed sequence of chromosome Y was not included in the reference (Rhie et al. 2023).

For each cell line, we calculated the total number of both unique and multi-mapped reads (Supplemental Table S2) and when all reads from the three cell lines were pooled, at least 10 reads were assigned to 40 out of 46 chromosome ends (19 short arms and 21 long arms; Fig. 1A and B; Supplemental Figs. S2-4; Supplemental Table S2). In the majority of the cases, read density peaked in the vicinity of the telomeric repeats and declined when moving inside the subtelomere, with reads terminating roughly 500 to 750 bp from the telomere (Fig. 1A and C; Supplemental Figs. S2-4). A striking different behavior was observed for the 18q and 22p (short arm) subtelomeres, where reads started 750 to 1000 bp away from the first telomeric repeat and covered a region of approximately 500 bp (Supplemental Figs. S2-4). This discrepancy may reflect differently structured 18q and 22p subtelomeres in the T2T genome and the cell lines analyzed in this study, and requires further clarification. Nonetheless, altogether these data reveal that: *i*) TERRA is transcribed from a large number of telomeres, as previously proposed (Azzalin et al. 2007; Nergadze et al. 2009; Arora et al. 2014; Episkopou et al. 2014; Porro et al. 2014; Feretzaki et al. 2019); and *ii*) most of TERRA molecules originate from regions very close to the first telomeric repeats and not from distant subtelomeric loci; this confirms that the heterogeneity of TERRA transcripts largely stems from the UUAGGG tracts (Azzalin et al. 2007; Farnung et al. 2012).

**Figure 1:**
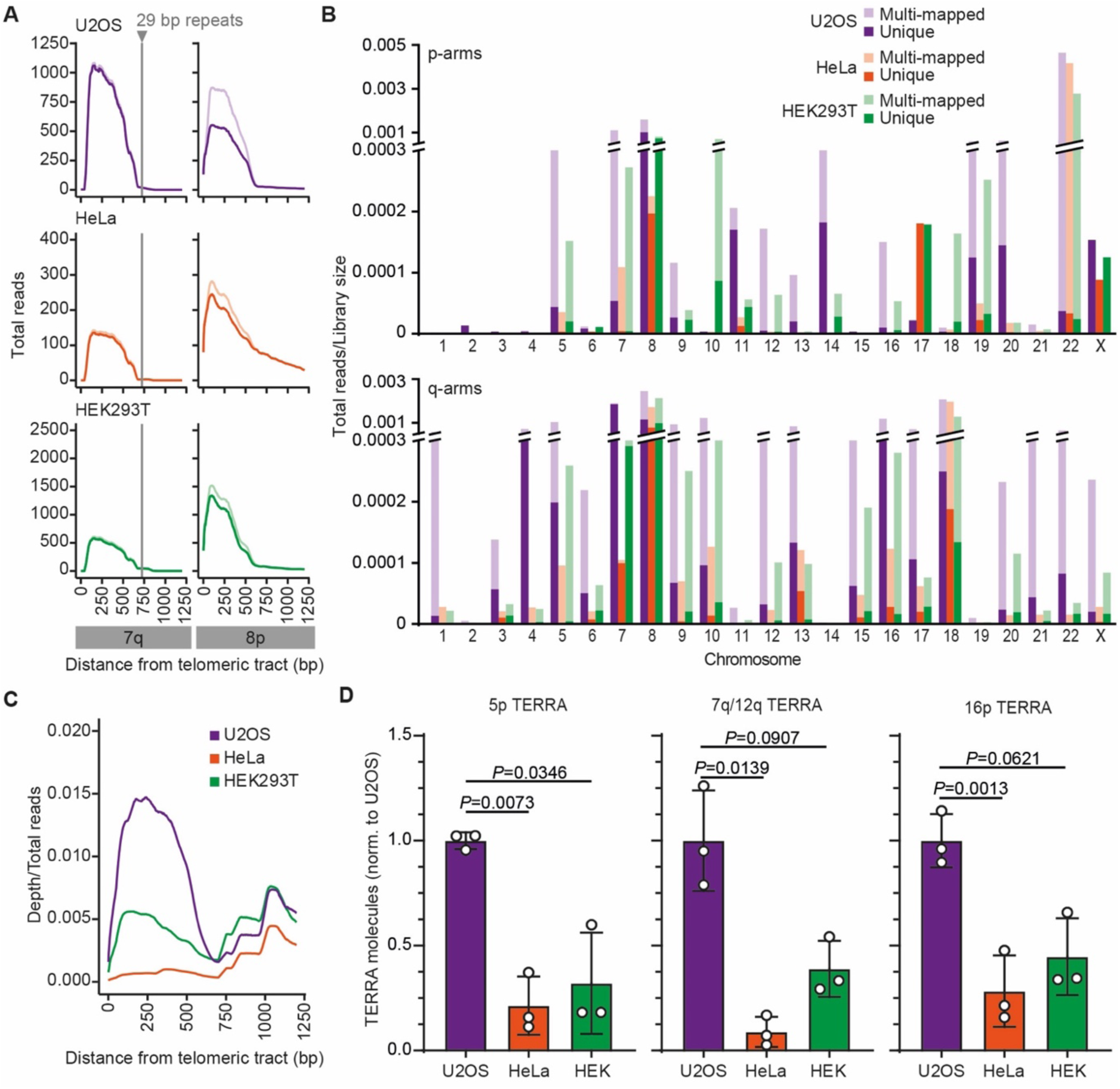
TERRA expression in different human cell lines. (A) TERRA ONTseq read coverage for 7q and 8p subtelomeres in U2OS, HeLa and HEK293T. Mapping was performed against the T2T CHM13v2.0/hs1 subtelomeric reference genome. Alignments span 1250 bp regions starting from the first telomeric repeat. The darker and lighter shade lines represent unique and multi-mapped reads, respectively. The vertical grey lines indicate the position of the most telomere proximal 29 bp-like repeat. (B) Quantification of the unique and multi-mapped reads (darker and lighter shades, respectively) within 1250 bp regions as in A for each human subtelomere. Reads are normalized to the corresponding library size. (C) Metaprofile of TERRA ONTseq of U2OS, HeLa and HEK293T cells. (D) RT-qPCR quantifications of TERRA transcripts from 5p, 7q/12q and 16p subtelomeres in the indicated cell lines. Values are normalized to the number of molecules in U2OS cells. Bars and error bars are means and SDs from 3 independent experiments. *P* values (Student’s *t* test) are indicated.

Read mapping across different chromosome ends was not uniform, with some ends being more frequently recognized. For example, the subtelomeres 8p, 7q and 8q were highly expressed in all cell lines (Fig. 1A and B; Supplemental Figs. S2-4; Supplemental Table S2). Conversely, some chromosome ends were under-represented in our alignments, including the ones of the short arms of the chromosomes 1, 2, 3, 4, 6, 15 and 21, and the long arms of chromosomes 2, 11, 14 and 19 (Fig. 1B; Supplemental Figs. S2-4; Supplemental Table S2). We conclude that TERRA transcripts from different ends are synthesized and/or degraded at different rates, as previously suggested based on the analysis of a few chromosome ends (Feretzaki et al. 2019; Savoca et al. 2023). Technical limitations, including differences in the efficiencies of purification (Supplemental Fig. S1C) and/or reverse transcription for different TERRA species, may have also contributed to the observed variabilities. Finally, we generated TERRA ONTseq metaprofiles and found that U2OS, HeLa and HEK293T showed similar profile shapes; however, U2OS cells had the highest overall read count, followed by HEK293T and HeLa (Fig. 1C). These data were validated using RT-qPCRs detecting a few selected TERRA species (Fig. 1D). Because ALT cells are characterized by elevated TERRA levels (Lovejoy et al. 2012; Arora et al. 2014; Episkopou et al. 2014), TERRA ONTseq appears to be able to quantitatively measure TERRA transcripts not only across different chromosome ends but also different samples.

### TERRA ONTseq identifies TERRA transcription start site regions

To locate TERRA promoters and transcription start sites (TSSs), we first re-analyzed the distribution of tandem repeats related to the canonical 29 bp repeats in our T2T-derived subtelomeric reference and identified them in almost all subtelomeres, with the exception of 6p, 8p, 13p, 14p, 17p, 21p, 22p, Xp, 8q, and 18q (Fig. 2; Supplemental Table S4). More specifically, we found 12 subtelomeres containing the canonical 29 bp sequence (Cons1, Fig. 2; Supplemental Table S4); 13 subtelomeres containing the canonical 29 bp sequence with a single mismatched nucleotide (Cons2); 8 subtelomeres containing a 29 bp sequence with 2 mismatches compared to Cons1 and 1 to Cons 2 (Cons2+1); and 8 subtelomeres containing a 29 bp sequence with a variable number of mismatches with respect to Cons1 (Cons1+n). In addition, the 9q subtelomere contains a 29 bp-repeat with two mismatches with respect to Cons2 (Cons2+2, Fig. 2; Supplemental Table S4); the 12q subtelomere contains a 30 bp sequence with 1 nucleotide insertion and 2 mismatches relative to Cons1 (Cons1+gap); and finally, the 1q and 17q subtelomeres contain a 34 bp-repeat sequence with two insertions of 2 and 3 nucleotides and one mismatch with respect to Cons2 (Cons3). We also predicted telomere-adjacent CpG islands using the T2T reference sequence and identified at least one of them within 2000 bp from the first telomeric repeat on all subtelomeres with the exception of the ones of 13p, 17p, 21p, 22p, Xp, and 8q arms. The identified CpG islands are comprised between 259 and 1553 bp (in 12q and 4p, respectively) and often contain multiple repetitions of one or more 29 bp consensus sequences (Fig. 2).

**Figure 2:**
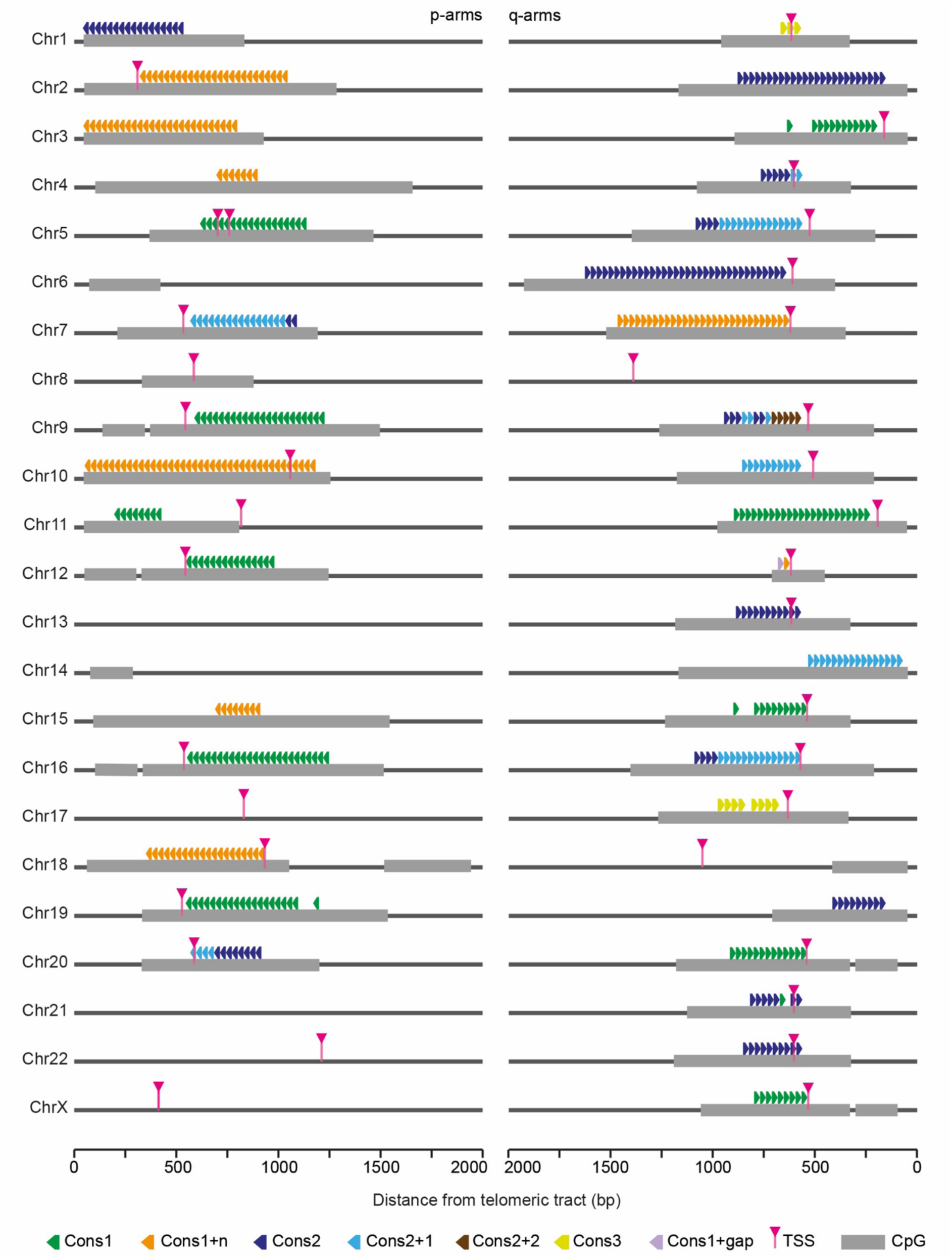
29 bp-like repeats, CpG-rich islands and TERRA TSSs across all human chromosome ends. Schematic representation of the positions of the different variants of 29 bp-like repeats (triangles) and subtelomeric CpG islands (grey boxes) in the T2T CHM13v2.0/hs1 subtelomeric reference genome. TERRA TSSs identified by TERRA ONTseq are also indicated (pink arrowheads). For each chromosome arm, the most telomere-proximal 2000 bp are shown. Note that chromosome 5p has two TERRA TSSs. The one on the left was identified in HeLa cells while the one on the right was identified in U2OS and HEK293T cells.

We then utilized TERRA ONTseq reads to identify TSS-containing regions by defining a threshold of at least 3 subtelomeric reads with 5’ ends mapping to the same 10 bp window. This strategy successfully identified 10 bp regions containing TERRA TSSs in 35 subtelomeres (Fig. 2; Supplemental Table S3). Notably, TERRA TSS regions are predominantly positioned within the CpG islands, just downstream of the 29 bp repeats (Fig. 2). This validates the ability of our pipeline to define TERRA TSSs and confirms that TERRA promoters are comprised of a variety of tandemly repeated 29 bp-like sequences. Importantly, some of the identified 29 bp-like repeats are only a few nucleotides away from the first telomeric repeats, as it is the case for the 1p, 3p, 10p and 14q subtelomeres (Fig. 2), for which virtually no unique TERRA reads were detected (Fig. 1B; Supplemental Figs. S2-4; Supplemental Table S2). It is plausible that those chromosome ends are also transcribed into TERRA molecules, but those transcripts escaped our analysis because the subtelomeric tract is too short to be mapped.

### TERRA ONTseq detects quantitative differences in TERRA molecules

As mentioned, TERRA ONTseq appears to be able to detect quantitative differences across samples different samples. To corroborate this assumption using isogenic cell systems, we took advantage of a previously described U2OS-derived cell line (SID4 cells) where TERRA expression from subtelomeres containing a 20 bp sequence within the 29 bp repeat consensus can be suppressed using activator-like effectors (T-TALEs) fused to the transcription repressor domain SID4X and cloned under the control of a doxycycline (dox) inducible promoter (Silva et al. 2021). Control cells expressing unfused T-TALEs (NLS3 cells; (Silva et al. 2021)) were also used. SID4 and NLS3 cells were treated with dox for 24 hours or left untreated and subjected to TERRA ONTseq (Supplemental Fig. S5A). T-TALE expression and suppression of TERRA transcription from selected chromosome ends containing 20 bp target sequences were validated by western blot and RT-qPCR, respectively (Supplemental Fig. S5B and C).

SID4 cells generated lower TERRA read counts (normalized to library size) than NLS3 cells already in absence of dox treatment, possibly due to leaky expression of the T-TALE constructs (Supplemental Fig. S5A and B). The dox treatment decreased the number of reads mapped to subtelomeres containing the T-TALE target in the SID4 samples (average 3.4-fold decrease), while a milder effect was observed at subtelomeres devoid of the 20 bp T-TALE target (average 1.4-fold decrease; Supplemental Fig. S5A; Supplemental Table S2). In NLS3 cells, dox treatment did not cause noticeable changes in TERRA reads at any chromosome ends (Supplemental Fig. S5A; Supplemental Table S2). Overall, these data confirm that TERRA ONTseq detects quantitative differences in TERRA levels between independent samples. Moreover, suppression of TERRA transcription from a fraction of chromosome ends does not trigger a compensatory mechanism increasing transcription from the other ends, at least in U2OS cells.

While analyzing the T-TALE data, we noticed that the 6q and 7q subtelomeres did not respond to dox treatment, despite being annotated in the T2T genome as containing numerous copies of the 20 bp target sequence (Supplemental Fig. S6A; Supplemental Table S4). This raised the possibility that the 6q and 7q subtelomeres contain 29 bp repeat-like sequences in the CHM13 cell line that was used for T2T sequencing (Nurk et al. 2022) but not in other cell lines. We thus focused on the highly expressed 7q subtelomere, which is annotated in the T2T genome assembly as having 29 copies of the Cons1+n 29bp repeat variant (Supplemental Fig. S6A; Supplemental Table S4). The same locus is devoid of 29 bp repeats in the GRCh38/hg38 human genome assembly (Supplemental Fig. S6A and B; Supplemental Table S4). Additionally, a previous study on the architecture of 7q subtelomere did not detect 29 bp repeats in 10 DNA samples from unrelated individuals (Baird et al. 2000). We PCR amplified HeLa and U2OS genomic DNA with oligonucleotides flanking the 29 bp repeats in the T2T 7q subtelomere and obtained only amplicons of sizes consistent with the lack of the 29 bp repeats tract (1537 and 697 bp for fragments with and without insertion, respectively; data not shown). We also cloned the amplified fragments into a plasmid and Sanger sequenced independent inserts for each cell line, and did not detect 29 bp repeat-like sequences in any of the inserts (Supplemental Fig. S6A and B). Hence, the 29 bp repeats present in the T2T genome at the 7q subtelomere might correspond to an insertion polymorphism that is not shared by all individuals or by HeLa and U2OS cells. This explains why our T-TALEs did not affect the expression of 7q TERRA and suggests a similar explanation also for the 6q subtelomere. Given these findings, we used the GRCh38/hg38 sequence for the 7q alignments performed later during this study.

### Analysis of 7q TERRA expression

The highly expressed 7q TERRA accounts for 26, 5 and 10% of total unique TERRA reads in U2OS, HeLa and HEK293T cells, respectively (Fig. 3A). To better characterize this transcript, we first performed northern blotting using a probe corresponding to a 441 bp fragment from the 7q subtelomere (7q/12q probe; Supplemental Fig. S6C). Due to the high identity between 7q and 12q subtelomeres (Supplemental Fig. S6C), our probe is expected to detect TERRA species from both loci; nonetheless, based on our TERRA ONTseq data, 74% of the signal generated by this probe should derive only from 7q TERRA (Fig. 1B; Supplemental Table S2). We isolated total and nuclear RNA from U2OS, HeLa and HEK293T cells and subjected it to northern blotting using 7q/12q and total (UUAGGG) TERRA probes. The signals generated by the two probes were similar and appeared as smears ranging from below 0.5 to more than 9 kb (Fig. 3B and C).

**Figure 3:**
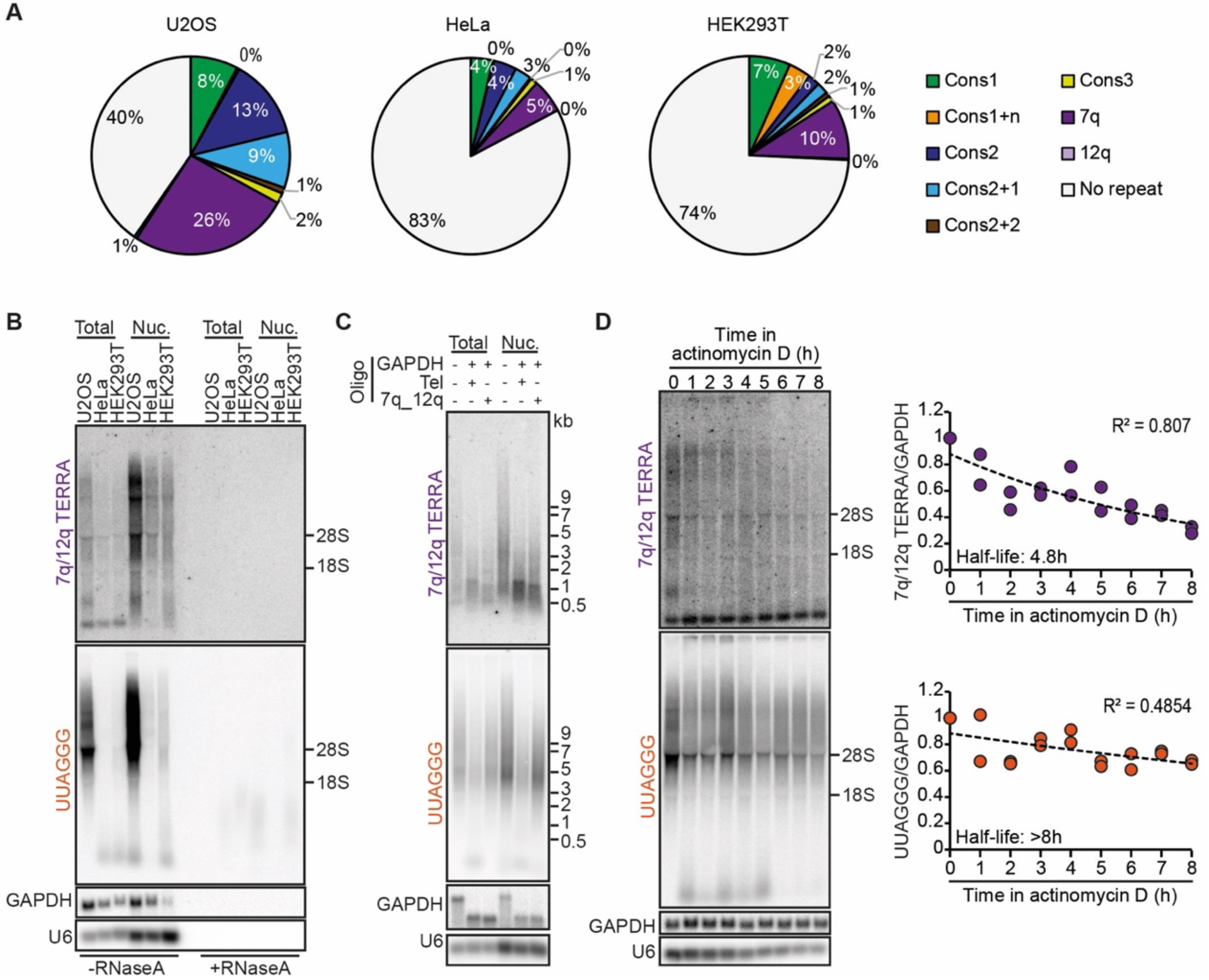
Expression of 7q TERRA. (A) Distribution of TERRA reads from 7q, 12q, 29 bp-like-containing subtelomeres and from repeat-devoid subtelomeres in the indicated cell lines. Only unique reads were considered for the analysis. (B) Northern blot hybridizations of total and nuclear RNA from U2OS, HeLa and HEK293T cells using probes for the 7q/12q TERRA or UUAGGG repeats (total TERRA). The positions of 28S and 18S ribosomal RNAs are indicated on the right. GAPDH and U6 were used as loading controls. RNaseA treatments were used to assure that signals derived from RNA and not genomic DNA contaminations. (C) Total and nuclear RNA from U2OS cells was incubated with RNaseH in the presence of oligonucleotides complementary to a sequence from 7q/12q TERRA or UUAGGG repeats (Tel). Oligonucleotides complementary to a GAPDH sequence were also included as indicated and served as a control for the RNaseH treatment. Products were analyzed by northern blotting as in B. (D) U2OS cells were treated with actinomycin D for the indicated times. Total RNA was extracted and analyzed by northern blotting as in C. The long lived GAPDH mRNA was used as a loading control. Half-lives and R^2^ values were calculated using a scatter plot analysis after the 7q/12q or the TERRA signals were normalized to the GAPDH signals. Data points are from 2 independent experiments and the values for time 0 are set to 1.

We then employed a northern blot-based approach previously developed to analyze TERRA transcripts from specific subtelomeres (Nergadze et al. 2009). We incubated total and nuclear RNA from U2OS cells with an oligonucleotide corresponding to a unique sequence shared by 7q and 12q subtelomeres and located 85 bp from the telomeric tract in the 7q subtelomere or with a (CCCTAA)5 oligonucleotide (7q_12q or Tel oligonucleotide, respectively; Supplemental Fig. S6C). In both reactions, we also included a control oligonucleotide annealing to GAPDH mRNA. We treated the RNA with RNaseH, which digests specifically RNA molecules engaged in DNA/RNA hybrids, and analyzed it by northern blotting (Fig. 3C). When compared to the undigested controls, the 7q/12q TERRA signal in RNaseH-treated sample incubated with the telomeric C-rich oligonucleotide appeared as a smear between approximately 1000 and 500 b; this smear should correspond to the 7q/12q TERRA subtelomeric sequences largely devoid of telomeric repeats. RNaseH digestion in presence of the 7q_12q oligonucleotide produced a slightly lower major band compared to the digestion with the telomeric oligonucleotide, confirming that the 7q/12q probe mainly detects 7q/12q TERRA and that these transcripts contain long telomeric UUAGGG tracts (Fig. 3C). Hybridization with a telomeric probe showed, as expected, a significantly reduced signal from the RNA incubated with the Tel oligonucleotide, while the incubation with the 7q_12q oligonucleotide led to a substantial disappearance of longer TERRA species above 9 kb (Fig. 3C). Attesting the efficiency and specificity of the RNaseH treatment, all GAPDH transcripts were shorter in samples treated with RNaseH while the length of the unrelated U6 transcript remained unaffected (Fig. 3C). Altogether, these experiments confirm that 7q TERRA constitutes a substantial fraction of total TERRA and reveals that individual 7q TERRA species contain long and heterogeneous UUAGGG repeat tracts in all tested cell lines.

To determine whether the increased levels of 7q TERRA derive from increased transcription or RNA stability, we measured 7q/12q TERRA stability by treating U2OS cells with actinomycin D to halt transcription and collecting total RNA every hour over 8 hours. Northern blot analysis showed that 7q/12q TERRA has a half-life of 4.8 hours while total TERRA has a half-life longer than 8h (Fig. 3D), which is consistent with previously reported values (Azzalin and Lingner 2008; Porro et al. 2010; Savoca et al. 2023). Hence the abundance of 7q TERRA cannot be ascribed to overall increased RNA stability, but rather it should derive from higher transcription rates.

### Analysis of 7q TERRA promoter regulation

7q/12q TERRA levels are higher in U2OS ALT cells than in HeLa and HEK293T telomerase positive cells, however those differences are not as important as the ones detected for total UUAGGG repeats (Fig. 3B). We expanded our analysis to a larger panel of cell lines and again observed that 7q/12q TERRA was, on average, 1.5 times higher in ALT cells compared to telomerase positive and primary lung fibroblasts, while the total UUAGGG signal was roughly 5.5 times higher in ALT cells compared to the others (Fig. 4A). We conclude that 7q/12q TERRA expression is not regulated identically to the majority of TERRA species.

**Figure 4:**
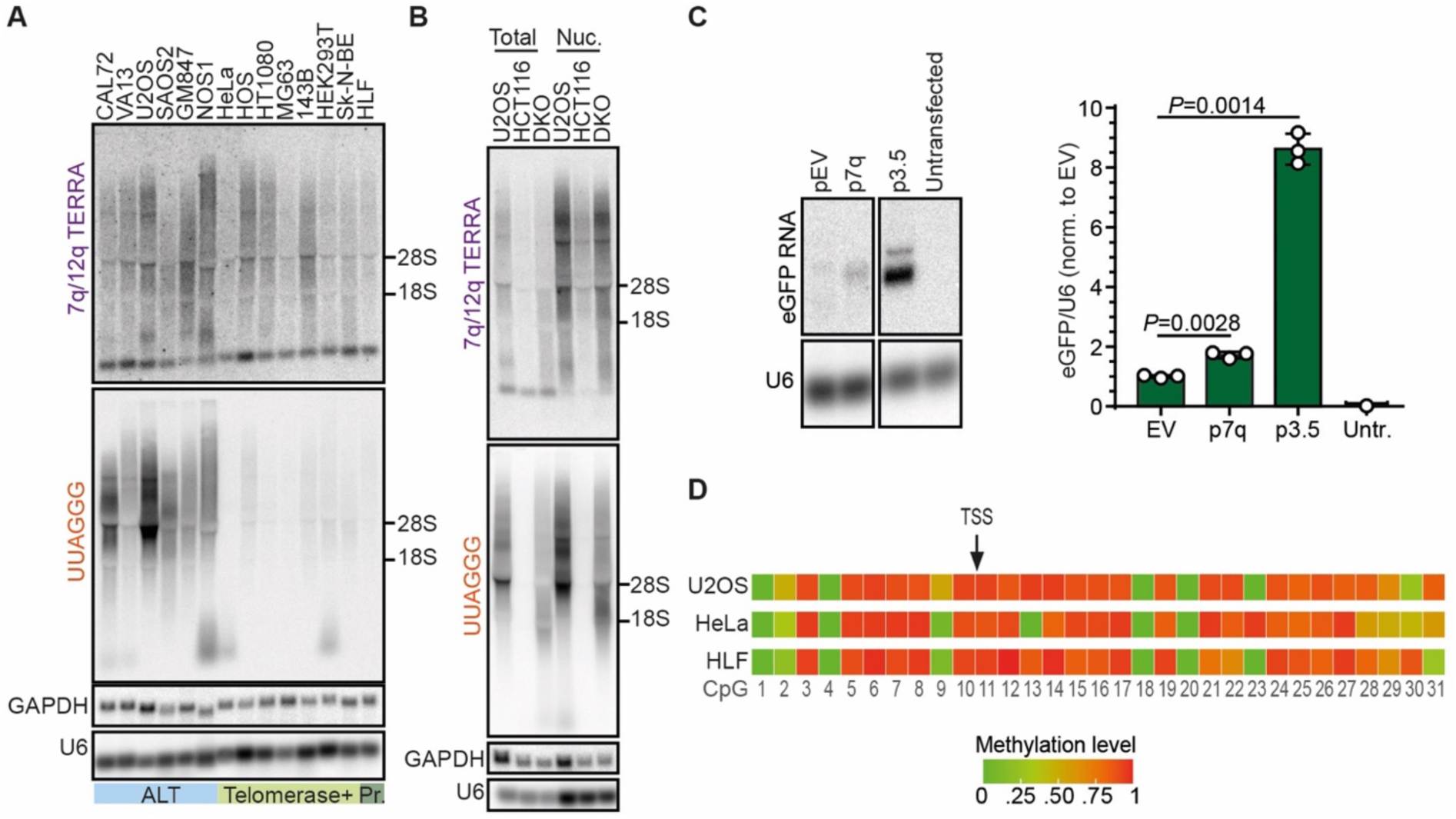
Transcriptional regulation of 7q TERRA. (A) Total RNA from ALT, telomerase positive (Telomerase+) or primary (Pr.) cells was analyzed by northern blotting using probes for the 7q/12q subtelomere or UUAGGG repeats (total TERRA). GAPDH and U6 RNAs were used as loading controls. (B) Total and nuclear RNA from HCT116 and DKO cells was analyzed by northern blotting as in A. U2OS RNA was included in the same blots for comparison with an ALT cell line. (C) Northern blot analysis of eGFP expression in HeLa cells transfected with the p7q promoter reporter. An empty vector plasmid devoid of promoter sequences (pEV) and the p3.5 plasmid containing a 29 bp repeat promoter were used as negative and positive controls, respectively. 24 hours after transfection, cells were selected using puromycin and 48 hours later harvested for RNA preparation. The graph shows the quantification of eGFP signals normalized through the corresponding U6 signals. Bars and error bars are the means and SDs from three independent experiments. *P* values (Student’s *t* test) are indicated. (D) Heatmap representation of 7q CpG island methylation for the indicated cell lines. The CpG island coordinates are chr7:159,335,203-159,335,443 in the GRCh38/hg38 reference genome. The position of 7q TERRA TSS gauged from TERRA ONTseq is indicated.

Because TERRA 29 bp repeats promoters are silenced through CpG methylation mediated by DNMT1 and DNMT3b (Nergadze et al. 2009; Feretzaki et al. 2019), it is possible that CpG methylation is not substantially affecting 7q/12q TERRA expression. Hence, we performed northern blots to analyze 7q/12q and total TERRA expression in human DKO cells and parental HCT116 cells. As expected, we detected a very faint signal for total TERRA in HCT116 parental cell lines when compared to the very strong signal detected for DKO cells (16-20 % of the DKO signal; Fig. 4B). On the contrary, 7q/12q TERRA signal was easily detected in HCT116 cells and its intensity was roughly half of the one in DKO cells (44-49% of the DKO signal; Fig. 4B). These data suggest that the 7q/12q promoters might be overall hypomethylated across different cell lines regardless of their telomere elongation mechanism, thus explaining why 7q/12q TERRA transcripts are more homogeneously expressed among cell lines and constitute a large fraction of total TERRA transcripts.

To test the hypothesis above, we first set up to isolate the 7q TERRA promoter. We generated a promoter reporter plasmid (p7q) by cloning a 695 bp fragment from the 7q subtelomere, overlapping a predicted CpG island of 333 bp (Fig. 2; Supplemental Fig. S6C), in front of an eGFP coding sequence. We transfected the plasmid into HeLa cells and analyzed eGFP expression by northern blotting. As a control, we transfected cells with a plasmid where the eGFP cDNA is preceded by a 29 bp repeat promoter from the XqYq subtelomere (p3.5 plasmid;(Nergadze et al. 2009)). We found that the selected 7q subtelomeric sequence was able to induce the expression of eGFP (Fig. 4C), confirming that we isolated a novel TERRA promoter sequence that does not contain 29 bp repeats. The p7q plasmid produced eGFP mRNA transcripts approximately 5 times less abundant than the ones produced from the p3.5 plasmid (Fig. 4C). This indicates that the 29 bp repeat promoters are stronger than the 7q promoter, however they are likely to be silenced more efficiently in cells through high-level CpG methylation.

Having defined the 7q promoter, we performed bisulfite sequencing of that region using genomic DNA from U2OS, HeLa, HEK293T, HLF, HCT116 and DKO cells and primers flanking the identified CpG island (Supplemental Fig. S6C). Unfortunately, we were not able to isolate amplification products for the 7q subtelomere from HEK293T, HCT116 and DKO cells, possibly due to polymorphisms within the regions annealing with the chosen oligonucleotides. Nonetheless, we could analyze CpG methylation in the other three cell lines. 7q methylation patterns from U2OS, HeLa and HLF did not show any striking differences (Fig. 4D; Supplemental Fig. S6C), indicating that CpG methylation has only a secondary role in regulating 7q TERRA expression when compared to 29 bp repeat promoters.

## CONCLUSIONS

TERRA ONTseq allows to simultaneously analyze TERRA transcripts from nearly all telomeres and provides robust and reproducible data. While confirming that 29 bp repeat-like sequences are main TERRA promoters, other promoter regions devoid of such repeats, such as the one of the 7q subtelomere identified here, are present in humans and substantially contribute to TERRA cellular levels. Similarly, TERRA ONTseq should allow to identify and validate all TERRA promoter sequences.

TERRA ONTseq also showed that the most transcribed chromosome ends are consistent between cell lines, likely pointing to the existence of conserved regulatory elements maintained across different cell types. Given the qualitative and quantitative nature of TERRA ONTseq, we are convinced that it will become an extremely useful tool for a rapid and precise analysis of the human TERRA transcriptome in different laboratorial and clinical settings. Further methodological improvements will broaden the reach of this tool, for example by developing protocols to also sequence the entire UUAGGG tract within TERRA or by adapting the TERRA ONTseq pipeline to organisms other than humans.

## MATERIALS AND METHODS

### Oligonucleotides and plasmid construction

Oligonucleotides were purchased from Integrated DNA Technologies and their sequences can be found in Supplemental Table S1. To generate the promoter reporter plasmid p7q, a 695 bp fragment from the 7q subtelomere was PCR amplified from HeLa genomic DNA using oligonucleotides #2384 and #2385 (Supplemental Table S1). The PCR product was cloned into the pJET1.2 plasmid using the CloneJET PCR Cloning Kit (Thermo Scientific), according to the manufactureŕs instructions. The resulting plasmid was digested with XhoI and NcoI and the insert was cloned in front of an eGFP cDNA. The p3.5 plasmid was a kind gift from Elena Giulotto (University of Pavia, Italy).

### Cell lines and tissue culture procedures

HeLa cervical cancer, HT1080 fibrosarcoma and HEK293T embryonic kidney cells were purchased from ATCC. U2OS osteosarcoma cells were a kind gift from Massimo Lopes (IMCR, Zurich, Switzerland). SAOS2, HOS, HOS 143B, a derivative of the HOS cell line, CAL72, NOS1 and MG63 osteosarcoma cells were a kind gift from Bruno Fuchs (Balgrist University Hospital, Zurich, Switzerland). Sk-N-BE neuroblastoma cells were a kind gift from Olga Shakhova (University Hospital Zurich, Zurich, Switzerland). VA13 lung and GM847 skin SV40-transformed fibroblasts were a kind gift from Arturo Londoño-Vallejo (CNRS, Paris, France). HLF primary lung fibroblasts were a kind gift from Joachim Lingner (EPFL, Lausanne, Switzerland). HCT116 human colon carcinoma cells and DKO cells were a kind gift from Bert Vogelstein, (Johns Hopkins University, Baltimore, USA). The LookOut Mycoplasma PCR Detection Kit (Sigma-Aldrich) was used to assure that all cell lines were mycoplasma-free. HeLa, HT1080, HEK293T, U2OS, HCT116, and DKO cells were cultured in high glucose DMEM, GlutaMAX (Thermo Fisher Scientific) supplemented with 5% tetracycline-free fetal bovine serum (FBS; Pan BioTech) and 100 U/ml penicillin-streptomycin (Thermo Fisher Scientific). WI-38 VA13 and HLF cells were cultured in high glucose DMEM, GlutaMAX supplemented with 10% tetracycline-free FBS and 100 U/ml penicillin-streptomycin. GM847 cells were cultured in high glucose DMEM, GlutaMAX supplemented with 10% tetracycline-free FBS, 100 U/ml penicillin-streptomycin and 1% nonessential amino acids (Thermo Fisher Scientific). NOS1 cells were cultured in high glucose RPMI 1640, GlutaMAX (Thermo Fisher Scientific), 5% tetracycline-free FBS, 1% sodium pyruvate (Thermo Fisher Scientific), and 100 U/ml penicillin-streptomycin. SAOS2, HOS, CAL72, MG63 and Sk-N-BE cells were cultured in high glucose DMEM/F12, GlutaMAX (Thermo Fisher Scientific), supplemented with 10% tetracycline-free FBS, 100 U/ml penicillin-streptomycin and 1% nonessential amino acids. 143B cells were cultured in high glucose DMEM/F12, GlutaMAX, supplemented with 10% tetracycline-free FBS and 100 U/ml penicillin-streptomycin. U2OS T-TALE cells were previously described (Silva et al. 2021) and were maintained in the same conditions as U2OS cells. For T-TALE expression, 100 ng/ml doxycycline (Sigma-Aldrich) was added to the culture medium for 24 h. For promoter reporter assays, HeLa cells were transfected using the Lipofectamine 2000 reagent (Invitrogen). Twenty-four hours after transfection, cells were incubated in medium containing 1 μg/ml puromycin (VWR International) and selected for 2 days.

### RNA preparation and analysis

RNA from whole cells or nuclei-enriched cellular fractions was prepared using the TRIzol reagent (Invitrogen) as previously described (Azzalin et al. 2007). RNA was treated twice with RNase-free DNase I (New England Biolabs) for northern blotting analysis and TERRA purification or three times for RT-qPCR analysis. For RNaseH experiments, 10 μg of nuclear RNA or 15 μg of total RNA were mixed with 625 nM GAPDH oligonucleotide (#134, Supplemental Table S1) and 625 nM telomeric oligonucleotide (#18, Supplemental Table S1) or 7q_12q oligonucleotide (#2336, Supplemental Table S1). The mix was incubated at 65°C for 4 min and then at room temperature for 20 min. 10 units of RNaseH (New England Biolabs) were added to the mix and allowed to digest at 37°C for 1 h. For northern blots, 10-15 μg of RNA were electrophoresed in 1 % formaldehyde agarose gels and blotted onto nylon membranes. Membranes were then hybridized for about 18 h in Church buffer containing ^32^P-labeled probes at 50°C-64°C. The strand-specific telomeric probe used to detect total TERRA was described previously (Azzalin et al. 2007). The 7q/12q probe was obtained by performing PCR on the plasmid p7q with the oligonucleotides #2384 and #1438 (Supplemental Table S1). The PCR product was strand-specifically radiolabeled by primer extension using the #2384 oligonucleotide. The eGFP probe was a DNA fragment obtained by digestion of the p3.5 plasmid with BamHI and XbaI and labeled by primer extension using the oligonucleotide #123 (Supplemental Table S1). The probes detecting β-actin, GAPDH and U6 RNAs were 5ʹ end-labeled DNA oligonucleotides (#5, #134 and #217, respectively; Supplemental Table S1). After hybridization, membranes were washed in 0.2-2× SSC, 0.2% SDS at the same temperatures used for hybridizations. Signals were detected using a Typhoon IP imager (Amersham) and quantified using ImageJ software. For quantitative RT-PCR experiments, 5 μg of total RNA or 1.5 μg of nuclear RNA were reverse transcribed with 0.5 μM TeloR and 0.05 μM U6 oligonucleotides (#18 and #217, respectively; Supplemental Table S1) using the SUPERSCRIPT IV reverse transcriptase (Invitrogen). Quantitative PCRs were performed with specific oligonucleotide pairs as indicated in Supplemental Table S1 using the iTaq Universal SYBR Green Supermix (BioRad) and a Rotor-Gene 6000 instrument (Corbett). Absolute TERRA quantifications were performed as previously described (Feretzaki et al. 2019) using plasmids containing subtelomeric fragments from specific chromosome ends to generate standard curves.

### DNA methylation analysis

Genomic DNA was extracted using the Promega Wizard Genomic DNA Purification Kit according to the manufactureŕs instructions. Bisulfite treatment was performed on 500 ng of DNA using the EZ DNA Methylation-Lightning Kit (Zymo research). Bisulfite-converted DNA was amplified using the oligonucleotides #2479 and #2480 (Supplemental Table S1) and the Q5U Hot Start High-Fidelity DNA Polymerase (New England Biolabs). PCR products were sequenced at Eurofins Scientific using the NGSelect Amplicon option (2-step amplicon generation workflow).

### Western blotting

Cells pellets were resuspended in lysis buffer (4% SDS, 20% Glycerol, 120 mM Tris-HCl pH 6.8), boiled at 95 °C for 5 min and centrifuged at 1600 × *g* at 4 °C for 10 min. Supernatants were recovered and protein concentrations determined using a NanoDrop 2000 (Tshermo Fisher Scientific). 35 μg of proteins were supplemented with 0.004% Bromophenol blue and 1% β-mercaptoethanol (Sigma-Aldrich), incubated at 95 °C for 5 min, separated in a 10% polyacrylamide gel, and transferred to a nitrocellulose membrane (Maine Manufacturing, LLC) using a Trans-Blot SD Semi-Dry Transfer Cell apparatus (Bio-Rad). Primary antibodies were a rabbit monoclonal anti-HA (Cell Signaling, #3724; 1:1000 dilution) and a mouse monoclonal anti-βActin (Abcam, ab8224; 1:5000 dilution). Secondary antibodies were HRP-conjugated goat anti-mouse and goat anti-rabbit IgGs (Bethyl Laboratories, A90-116P and A120-101P; 1:2000 dilution). Signal detection was performed using the ECL detection reagents (GE Healthcare) and an Amersham 680 blot and gel Imager.

### TERRA purification and ONT Sequencing

For each TERRA purification reaction, 25 µgs of nuclear RNA were incubated at 75°C for 4 min in 0.5X TE (5 mM Tris–HCl pH 7.5, 0.5 mM EDTA). 25 pmols of biotinylated, 3’ dideoxycytosine (ddC)-blocked telomeric oligonucleotides (#1, Supplemental Table S1) were added to the RNA and the mixture incubated at 50°C for 50 min in 0.5X TE supplemented with 1 M of NaCl. 50 μg of streptavidin magnetic beads (New England Biolabs) were added and the mixture was incubated in a thermomixer at 50°C with 300 rpm agitation for 3 h. Beads were washed once in 5 mM Tris–HCl pH 7.5, 0.5 mM EDTA, 1 M NaCl, 3 times in 5 mM Tris–HCl pH 7.5, 0.5 mM EDTA, 0.5 M NaCl and twice in 5 mM Tris–HCl pH 7.5, 0.5 mM EDTA, 0.15 M NaCl. RNA was eluted in 10 mM Tris–HCl pH 8.0 by heating the mixture for 3 min at 80°C. RNA was then treated with DNAseI at 37°C for 30 min and precipitated in the presence of glycogen. TERRA sequencing was performed using the Direct cDNA Sequencing Kit from Oxford Nanopore Technologies (SQK-DCS109) according to the manufactureŕs instruction with minor modifications during the cDNA synthesis. Specifically, 400-800 ng of purified TERRA, 10 nM of VNP-TERRA oligonucleotide (#2161, Supplemental Table S1) and 10 µM of dNTPs were mixed together and heated to 85°C for 3 min. The cDNA synthesis reaction was performed with SUPERSCRIPT IV reverse transcriptase (Invitrogen) at 55°C for 70 min followed by 20 min at 42°C. HeLa, HEK293T and T-TALE cDNAs were barcoded using the Native Barcoding Expansion kit from Oxford Nanopore Technologies (EXP-NBD104). Sequencing was performed using the MinION device with the flow cells MK 1 R9 version (Oxford Nanopore Technologies). For U2OS, a total of 225 µgs of nuclear RNA was used for 2 sequencing runs; for HeLa, 727 µgs of nuclear RNA was used for 6 sequencing runs (3 of which barcoded); for HEK293T 1 mg of nuclear RNA was used for 7 sequencing runs (4 of which barcoded); for T-TALES, 225 µgs of nuclear RNA was used for 3 sequencing barcoded runs.

### Nanopore sequence analysis

Nanopore reads were de-barcoded using the Porechop tool (https://github.com/rrwick/Porechop). The alignment to the T2T CHM13v2.0/hs1 genome reference or a derived subtelomeric reference was performed using the minimap2 aligner (Li 2018). The options used for the alignment step were *-k15 – w6* (minimizers on the query), *-ax map-ont* (for Oxford nanopore reads). The alignment results were statistically analyzed through the functions *stats* and *idxstats* from samtools (Danecek et al. 2021). The mapping quality is given by the MAPQ value, that describes the uniqueness of the alignment. MAPQ=0 indicates the reads that can possibly map to many locations and have the lowest quality, while MAPQ> 0 are primary reads, or uniquely mapped. The coverage of the aligned reads was calculated using the function *genomeCoverageBed* from bedtools suite (Quinlan and Hall 2010). Multi-mapped reads where distributed equally along the reference sequence. Custom scripts in R language scripts were set up for coverage visualization, comparison among the different cell lines and for the position of 29 bp and 20 bp repeats.

### DNA and bisulfite sequencing analysis

7q and 12q subtelomere sequences were aligned using the EMBOSS Needle tool at the EMBL’s European Bioinformatics Institute (https://www.ebi.ac.uk/Tools/psa/emboss_needle/). CpG dinucleotide contents were analyzed using the CpGPlot software (https://www.ebi.ac.uk/Tools/seqstats/emboss_cpgplot/). The distribution of 29 bp repeat-like sequences was determined using the simple repeat annotation and BLAT at UCSC Genome Browser (https://genome.ucsc.edu/). For methylation analysis, the forward and reverse reads from bisulfite sequencing were merged using the *fastq-join* function (https://manpages.ubuntu.com/manpages/bionic/man1/fastq-join.1.html#name) and the *filter-fastq* function (https://github.com/Floor-Lab/filter-fastq) was used to select reads shorter than 400 bp to perform the mapping to 7q and reads longer than 400 bp to 12q. Read mapping was performed using the software *bwameth* (https://github.com/brentp/bwa-meth) using a FASTA reference containing the 7q and 12q subtelomeric sequences. The computation of methylation metrics from alignment files was performed with *MethylDackel* software (https://github.com/dpryan79/MethylDackel). A custom script in R language was set up to generate methylation state heatmaps.

### Statistical analysis

For direct comparison of two groups, a paired two-tailed Student’s *t* test was performed using Microsoft Excel. *P* values are shown in the figures.

## Data availability

TERRA ONTseq data were deposited at the Sequence Read Archive (SRA) with the code PRJNA997813. Sanger and bisulfite sequencing data can be provided upon request.

## ACKNOWLEDGMENTS

We thank Miguel Alves for help with isolating and sequencing the 7q subtelomere, Patricia Abreu for critical reading of this manuscript and the Bioimaging and Information System units of iMM for services. We also thank Elena Giulotto, Massimo Lopes, Bruno Fuchs, Olga Shakhova, Arturo Londoño-Vallejo, Joachim Lingner, and Bert Vogelstein for sharing reagents. This work was supported by LaCaixa Foundation (project LCF/PR/HP21/52310016 to C.M.A.), national funds through FCT – Fundação para a Ciência e a Tecnologia, I.P. (projects PTDC/BIA-MOL/6624/2020 and 2021.00143.CEECIND to C.M.A), TessellateBIO (to C.M.A) and CNR – Consiglio Nazionale delle Ricerche (project DBA.AD005.225 - NUTRAGE-FOE2021 to S.B.).

## AUTHOR CONTRIBUTIONS

J.R. and C.M.A. conceptualized the study and designed the experiments. J.R. performed the experiments. R.A. and S.B. performed all bioinformatic analyses. J.R. and C.M.A. wrote the manuscript and all authors reviewed and edited the manuscript. C.M.A. and S.B. acquired funding and provided supervision.

## Supplemental information

### SUPPLEMENTAL FIGURES

**Supplemental Figure S1:**
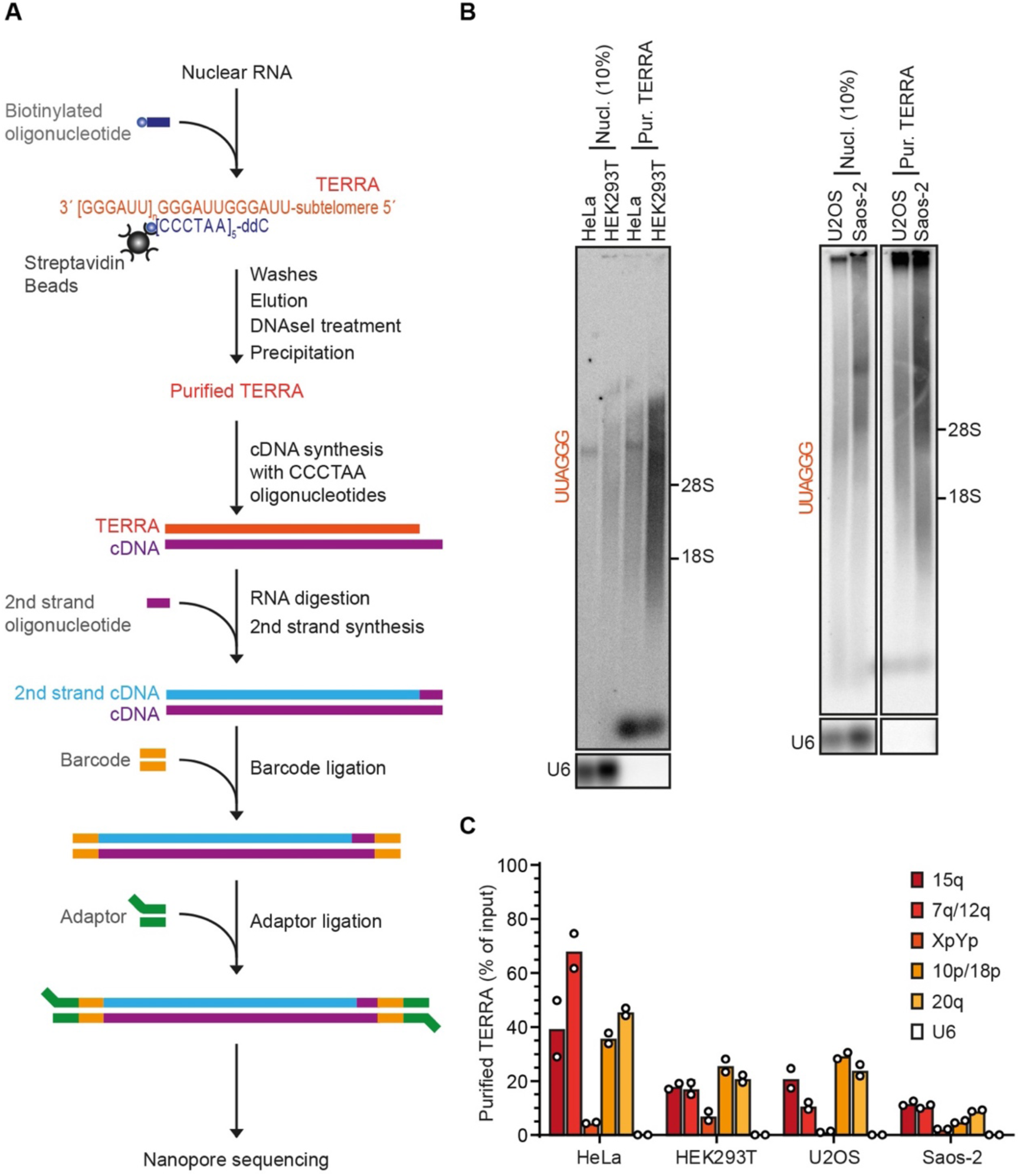
TERRA purification and sequencing. (A) Schematic representation of the TERRA purification and ONT sequencing pipeline. (B) TERRA-enriched nuclear RNA from the indicated cell lines was analyzed by northern blotting using a probe complementary to the UUAGGG repeats (total TERRA) or U6. (C) RT-qPCR quantifications of TERRA transcripts or U6 in TERRA-enriched nuclear RNA as in B. Values are expressed as fraction of the input. Bars are the averages from two independent experiments.

**Supplemental Figure S2:**
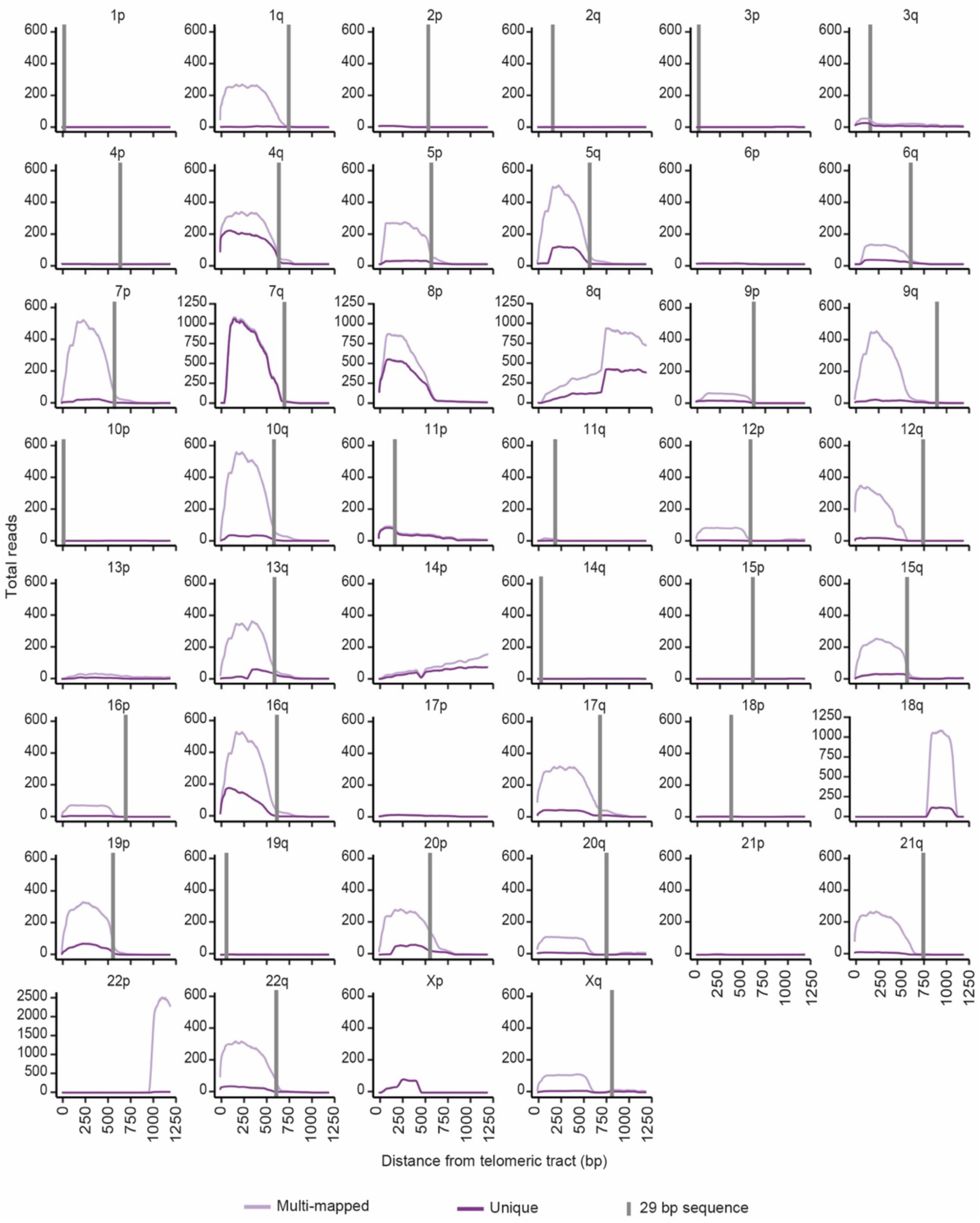
TERRA ONTseq mapping for U2OS cells. ONT reads from U2OS cells mapped to the T2T CHM13v2.0/hs1 subtelomeric reference genome. Read coverage results for the most telomere-proximal 1250 bp of subtelomeres are shown. Light and dark purple lines represent multi-mapped and unique reads, respectively. Vertical grey lines indicate the position of the most telomere-proximal 29 bp-like repeat.

**Supplemental Figure S3:**
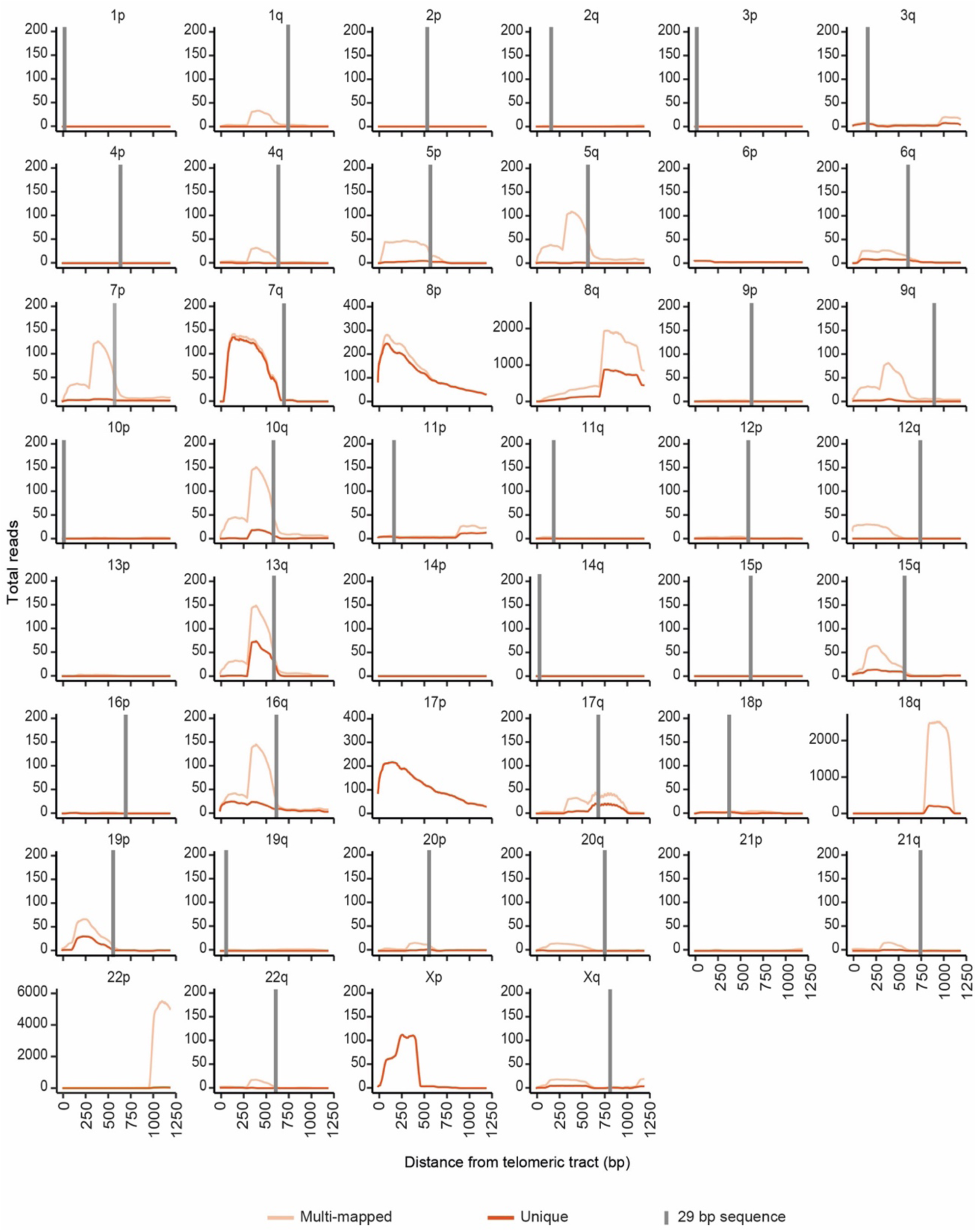
TERRA ONTseq mapping for HeLa cells. ONT reads from HeLa cells mapped to the T2T CHM13v2.0/hs1 subtelomeric reference genome. Read coverage results for the most telomere-proximal 1250 bp of subtelomeres are shown. Light and dark orange lines represent multi-mapped and unique reads, respectively. Vertical grey lines indicate the position of the most telomere-proximal 29 bp-like repeat.

**Supplemental Figure S4:**
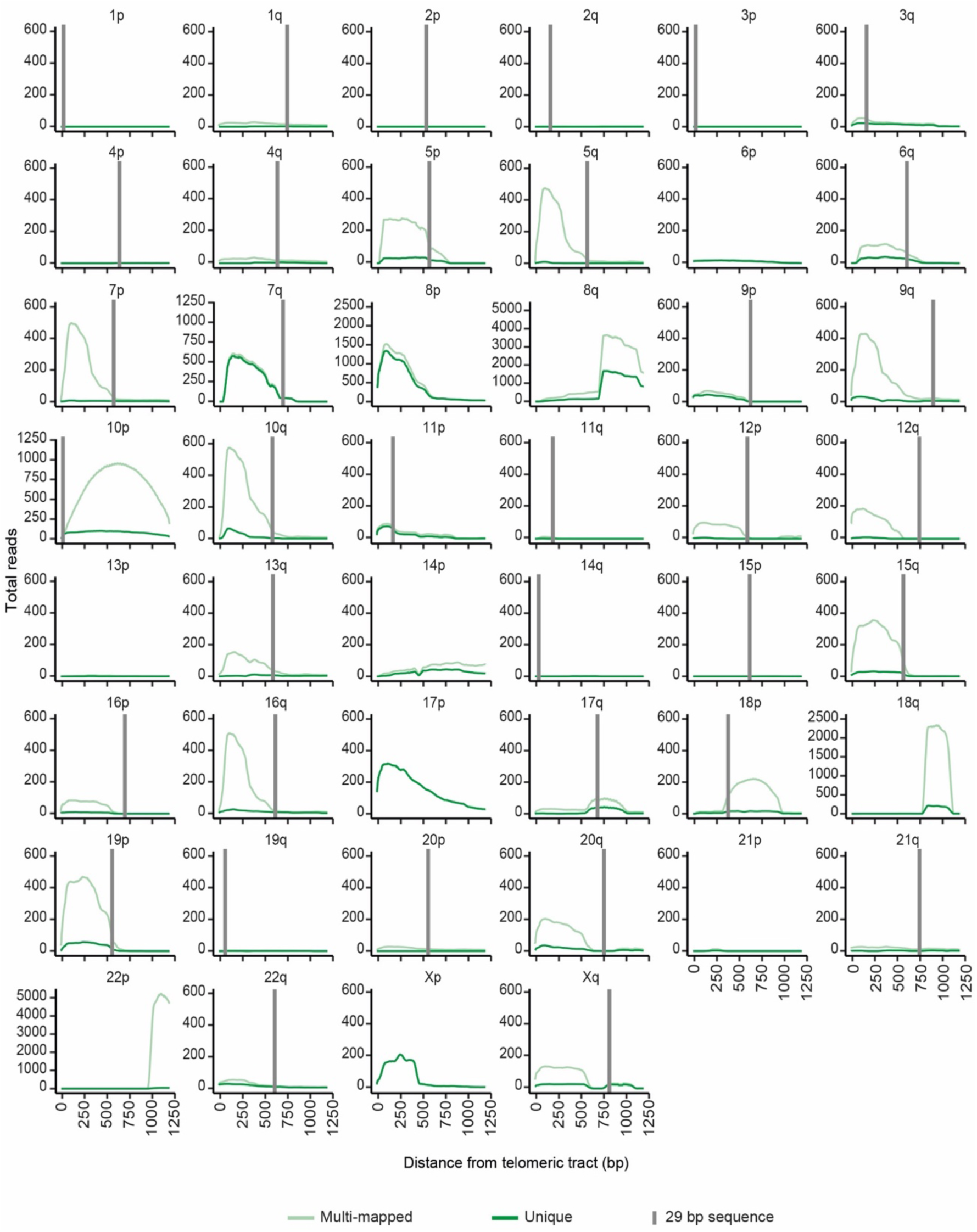
HEK293T TERRA ONTseq mapping for HEK293T cells. ONT reads from HEK293T cells mapped to the T2T CHM13v2.0/hs1 subtelomeric reference genome. Read coverage results for the most telomere-proximal 1250 bp of subtelomeres are shown. Light and dark green lines represent multi-mapped and unique reads, respectively. Vertical grey lines indicate the position of the most telomere-proximal 29 bp-like repeat.

**Supplemental Figure S5:**
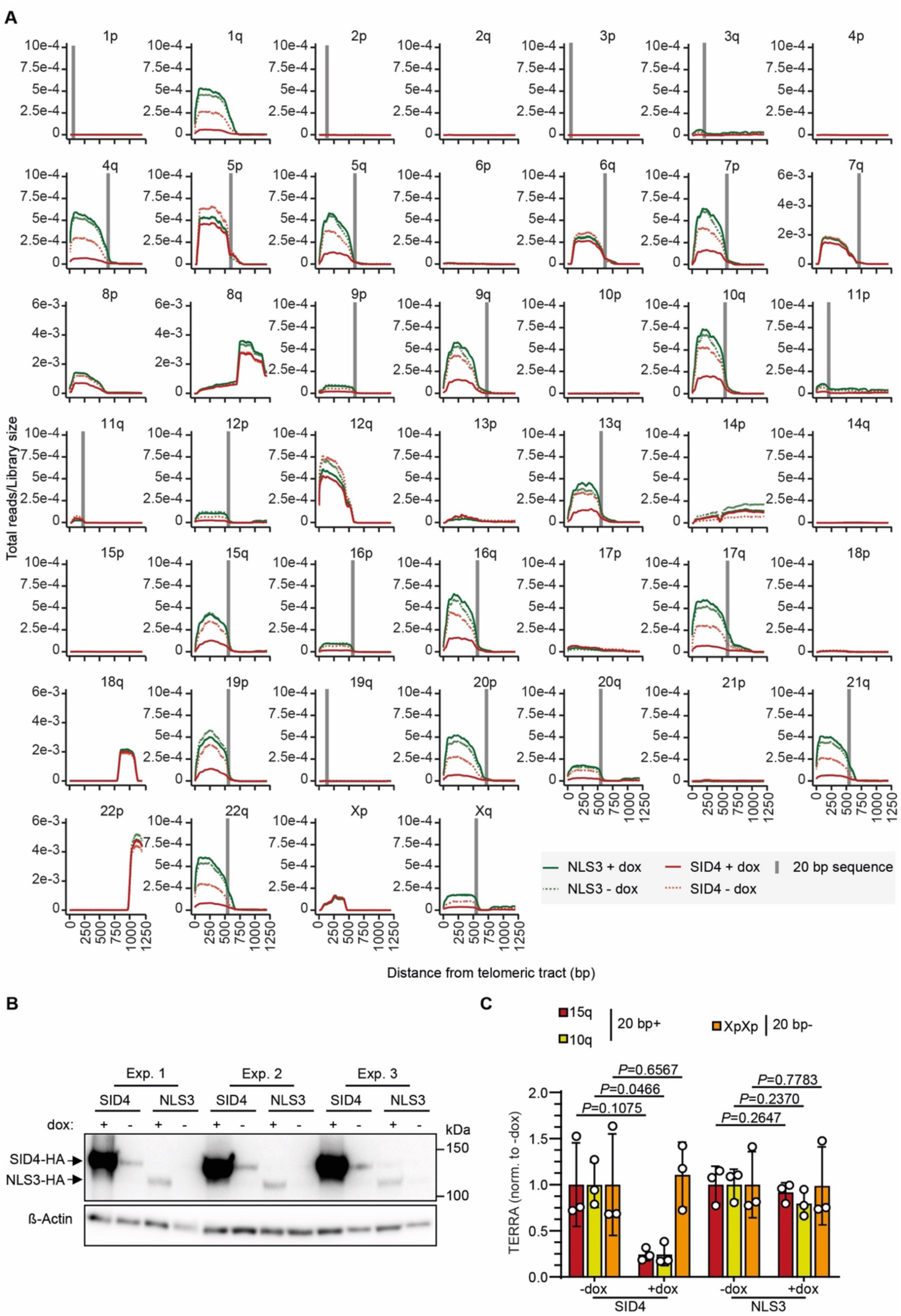
TERRA ONTseq mapping for T-TALE cell lines. (A) ONT reads from T-TALE cells mapped to the T2T CHM13v2.0/hs1 subtelomeric reference genome. Read coverage results for the most telomere-proximal 1250 bp of subtelomeres are shown. Green lines represent the reads from NLS3 cells treated with dox for 24 h and dotted light green lines from untreated NLS3 cells; red lines represent the reads from SID4 cells treated with dox and dotted light red lines from untreated SID4 cells. Vertical grey lines indicate the position of the most telomere-proximal 20 bp T-TALE target (100% identity), which is contained within the 29 bp-like repeat TERRA promoters. (B) Western blot analysis of total protein extracts from the same cells as in A using anti-HA antibodies to detect the HA tag fused to the N-terminus of the T-TALEs. Eta (β) Actin serves as loading control. (C) RT-qPCR quantifications of TERRA transcripts from subtelomeres containing (20 bp+; target) or devoid of (20 bp−; non-target) the 20 bp sequence recognized by the T-TALEs in cells as in A. After normalization through U6, values from dox-treated samples (+dox) were expressed as fold change over untreated (-dox). Bars and error bars are averages and SDs from three independent experiments. P values (Student’s t test) are indicated.

**Supplemental Figure S6:**
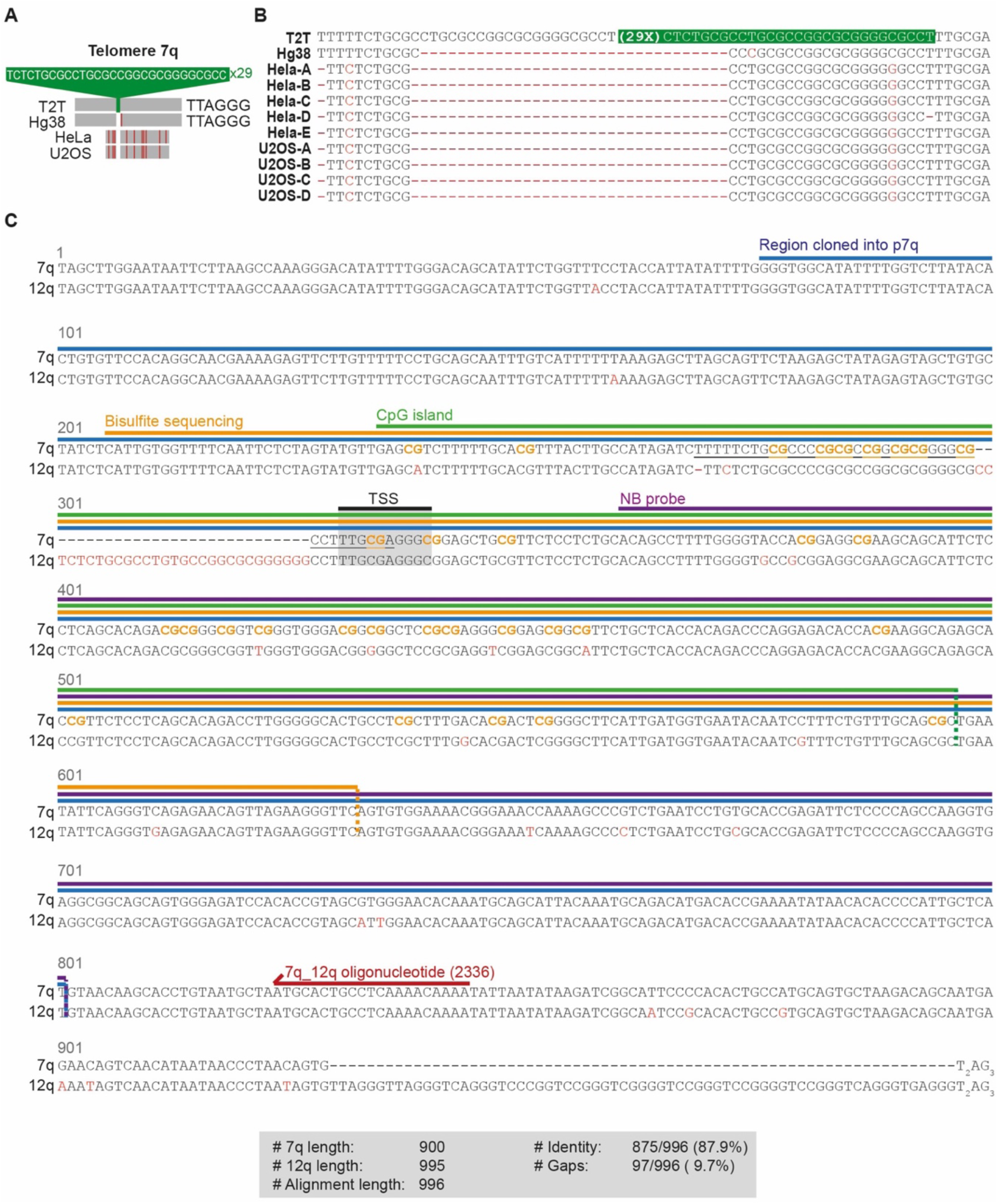
Genomic structure of the 7q subtelomere. (A) Schematic representation of 7q subtelomere regions from the T2T reference genome (coordinates: chr7:160,563,685-160,564,877), the corresponding region from the GRCh38/hg38 reference genome (coordinates: chr7:159,335,185-159,335,537) and HeLa and U2OS genomic DNA sequenced in this study. The DNA insertion in the T2T is in green and the single mismatches to the T2T sequence are in red. (B) Sequence alignments between a 7q subtelomeric fragment from the T2T reference genome, the same region from the GRCh38/hg38 reference genome and multiple PCR products obtained from HeLa and U2OS genomic DNA. Mismatches to the T2T sequence are in red. The sequence repeated in tandem and present only in the T2T reference sequence is boxed in green. (C) Sequence alignment between a 900 bp long fragment from the subtelomere of chromosome 7q (coordinates chr7:159,334,969-159,335,868 in the GRCh38/hg38 reference genome) and a highly similar sequence from the subtelomere of chromosome 12q (coordinates chr12:133,263,945-133,264,939 in the GRCh38/hg38 reference genome). Key features are indicated and include the fragment cloned into the p7q reporter plasmid (in blue), the fragment amplified for bisulfite sequencing (orange), the predicted CpG island (green), the TSS identified by TERRA ONTseq (black), the fragment corresponding to the probe used for northern blotting (purple), the 7q_12q oligonucleotide used in RNAseH experiments (red), and the region aligned in B (underlined). The 31 CpG dinucleotides present in the predicted CpG island are in orange.

### SUPPLEMENTAL TABLE LEGENDS

**Supplemental Table S1**. List of primers used in this study.

**Supplemental Table S2**. Read counts for TERRA ONTseq from U2OS, HeLa, HEK293T, U2OS SID4 (plus and minus dox) and U2OS NLS3 (plus and minus dox) cells.

**Supplemental Table S3**. 10 bp regions containing putative TERRA transcription start sites derived from TERRA ONTseq.

**Supplemental Table S4**. List of the different variants of the 29 bp-repeats and comparison of the 29 bp-promoter positions in the GRCh38/hg38 and the T2T CHM13v2.0/hs1 reference genomes.

